# Tactile and pain mechanical sensitivity of the human hand

**DOI:** 10.64898/2026.02.23.707479

**Authors:** María José Giner, Michael Mazar, Fernando Aleixandre-Carrera, Karel Talavera, Miguel Delicado-Miralles, Vicente Miralles-Liborio, Enrique Velasco

## Abstract

The human hand has a refined mechanical sensitivity, allowing it to play crucial roles in tactile exploration and object manipulation. Despite its fundamental and clinical relevance, a comprehensive characterization of mechanical sensitivity across the human palm is still lacking.

Here, we mapped the spatial distribution of innocuous and noxious mechanical sensitivity across the palmar surface of the human hand. We examined 66 hands from 33 healthy adults, dividing the palm into 27 areas, in each of which we measured the mechanical detection threshold, the mechanical pain threshold and the pain intensity evoked by a standard 300 g pinprick stimulus.

We found distal areas (i.e., fingertips) to exhibit higher tactile sensitivity than proximal areas (i.e., the wrist). Notably, the sensitivity to innocuous and noxious mechanical stimuli were inversely correlated across areas, such that areas with higher tactile sensitivity displayed higher pain thresholds. In addition, the dominant hand was less sensitive than the non-dominant one, and women displayed higher sensitivity than men.

Together, this work provides the first detailed spatial characterization of mechanical sensitivity across the human hand and introduces a systematic methodology for its assessment. These findings set the stage for future studies of the neurophysiological mechanisms of touch and pain in the human hand and for clinical research into pathological conditions involving the altered hand sensitivity.

## INTRODUCTION

The hand is our primary tactile sensor and object manipulator. The freedom of movement that arose with bipedalism and the parallel evolution of the human brain resulted in the hand acquiring a pivotal role in social interaction, skilled behavior, and cultural development (Napier, 1962). The hand interfaces with the external world through the skin, a complex sensory organ that can be broadly divided into glabrous (non-hairy; palms and soles) and hairy (covering the rest of the body) (Ackerley *et al*., 2014). In particular, the glabrous skin on the palm of the hand provides an indispensable source of tactile information. Mechanical forces are detected and transduced by mechanoreceptors: specialized cutaneous somatosensory receptors that encode texture, vibration, pressure and noxious mechanical stimuli (Johansson & Vallbo, 1979; Abraira & Ginty, 2013; Cobo *et al*., 2021).

The distribution of tactile sensitivity across the palmar surface has long been a subject of interest, with evidence suggesting it may not be uniform (Johansson & Vallbo, 1979). Given their implication in object exploration and manipulation, one could hypothesize that tactile sensitivity is highest at the fingertips, enabling precise and rapid tactile discrimination (Corniani & Saal, 2020; Jarocka *et al*., 2021). Hand sensitivity is frequently compromised in a wide range of conditions, including peripheral nerve injuries such as diabetic polyneuropathy and carpal tunnel syndrome, as well as neurological, muscular and joint diseases, and during aging (Novak *et al*., 1992; Farrell *et al*., 2000; Kalisch *et al*., 2008; Catley *et al*., 2014; Venkatesan *et al*., 2015; McIntyre *et al*., 2021). Consequently, the mechanical sensitivity of the hand has been extensively assessed in clinical studies (Gellis & Pool, 1977; Cole *et al*., 1998; Tremblay *et al*., 2005; Kalisch *et al*., 2008; Bowden & McNulty, 2013*a*), but despite its fundamental and clinical relevance, a detailed characterization of mechanical sensitivity across the human palm is still lacking. Here, we address this gap by systematically mapping the mechanical detection and pain thresholds along 27 areas of the palmar surface of the hand, and by examining the influence of sex, laterality and hand dominance.

## MATERIALS AND METHODS

### Subjects, ethics, and exclusion criteria

After obtaining approval from the Ethical Committee (Ethical Committee of pharmacological research in the University General Hospital of Elche, Alicante, Spain) a total of 33 healthy voluntaries were recruited (Fig. 1 and Table 1). Three subjects were excluded from the analysis for pain variables as none of the von Frey filaments elicited pain on them. The study was executed in the physical therapy clinic “Francisco Javier Ortega Physical Therapy”, located in Elche, Alicante, Spain.

**Table 1.** Descriptive data from the recruited sample (n = 33). “Drug consumption” refers to antihistamines, bronchodilators, or contraceptives. “Upper limb pathological history” includes the previous conditions elbow fracture, 5^th^ finger fracture, and 2^nd^ metacarpal fracture. “Physical activity” is a qualitative assessment based on whether the participant self-identifies as an athlete. “Toxic habits” refers to smoking or alcohol consumption.

| Characteristic | Value |
| --- | --- |
| Age (Mean ± SD) | 26.3 ± 2.1 |
| Woman / Man (n) | 15 / 18 |
| Right-handed / Left-handed (n) | 26 / 7 |
| Upper limb pathological history (n) | 3 |
| Physical activity (n) | 24 |
| Toxic habits (n) | 19 |
| Drugs consumption (n) | 5 |

**Figure 1.**
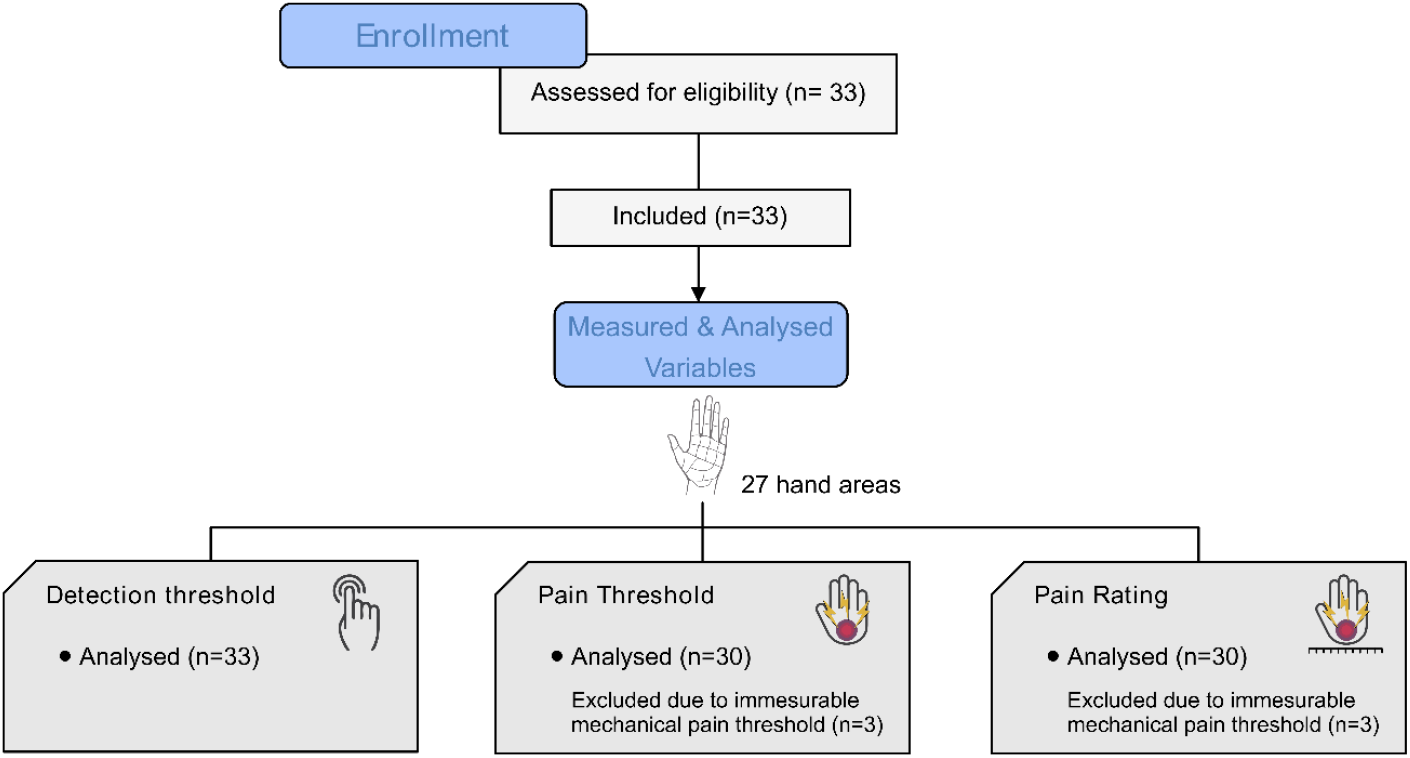
Experimental design and measured variables.

Exclusion criteria were as follows:

- Upper limb pathology within 30 days prior to the study.
- Severe systemic conditions, including diabetes mellitus, cancer, neurological disease, depression, fibromyalgia, and immunosuppression.
- Dermatological pathologies or skin alterations (e.g., tattoos or scars) at the testing sites.
- Use of anticoagulants, antidepressants, gabapentinoids, or opioids during the study or within the week prior to participation.
- Use of NSAIDs within 48 h of or during the study.
- Pregnancy.

Subjects gave written informed consent prior to any intervention, according to Helsinki’s and World Medical Association declarations, and completed a demographic questionnaire.

### Experimental design

The present study is a cross-sectional observational study, aiming to map the sensitivity of the palm of the hand towards innocuous and noxious mechanical stimuli in healthy patients.

Given the absence of an existing standardized methodology for assessing this, we established a novel method to uniformly characterize hand sensitivity. This involved measuring tactile and painful sensitivity at 27 different areas on the palmar surface (comprehending proximal, middle and distal phalanx; metacarpal heads; midpalmar line; thenar and hypothenar eminences; the midpoint between both eminences; the wrist) of both hands. These areas were grouped into six clusters (Fig. 1). To minimize bias, a pseudorandom number generator determined the order of hands (left or right), clusters, and the starting area for each cluster.

### Sensory evaluation

Participants were seated with their forearms supinated and relaxed. Vision was occluded by a mask to avoid visual detection of the stimuli. The variables assessed were (Fig. 1):

- *Mechanical detection thresholds* (hereafter, detection thresholds) were determined by applying von Frey filaments (Bioseb, bio-vf-m, Vitrolles) of increasing pressure five times over the skin and asking the subjects to inform when they felt the slightest touch from the filament. The threshold was defined as the first filament producing three successful reports (Beltrá *et al*., 2022).
- *Mechanical pain thresholds* (hereafter, pain thresholds) were determined using a similar procedure to the detection threshold but asking the subjects to inform when they felt minimum pain instead.
- *Evoked pain:* to evaluate the amount of pinprick pain elicited by a painful stimulus, we applied a standardized 300 g von Frey filament and asked the subjects to the pain evoked by it using a Verbal Rate Scale (VRS) (Chien *et al*., 2013), from 0 (no pain) to 10 (unbearable pain). The stimulus was applied twice, and the average of both ratings was calculated for analysis.

### Data analysis

#### Sample size calculation

The sample size calculation was based on a previous experimental trial from our group, in which we analyzed the same variables in healthy subjects at four different locations in the hand (Beltrá *et al*., 2022). For an alpha of 5% and a statistical power of 80%, the required sample size was 27 subjects. Finally, 33 subjects were recruited.

#### Statistical analysis

MS Excel (Microsoft Co., 2019) and Jamovi v.2.3 (The jamovi project, 2024) were used for data collection and statistical analysis, respectively. First, descriptive statistical analysis was performed to detect possible outliers and to determine the distribution of the variables. Subsequently, inferential statistics were conducted using mixed models. To investigate potential differences due to sex or hand dominance, these were included as fixed effects in the models for all the three variables, and the subject was included as a random effect to account for the repeated measures design across hand areas. On the one hand, a generalized mixed model (GMM) with a gamma distribution and a log-link function was used to analyze the detection thresholds. This model provided the best fit for this highly skewed data distribution, partially attributed to the inherent limitations of von Frey filaments in accurately determining very low detection thresholds as they approach 0. On the other hand, linear mixed models (LMMs) were used to analyze pain thresholds and evoked pain. To approximate a normal distribution of the data, pain thresholds were log-transformed. If differences were found in the models, a *post hoc* test was conducted, applying the Holm correction for multiple comparisons.

Results from the models are reported as estimated marginal means (EMMs), which are adjusted for the other factors in each model to provide a more precise estimate of the effect of interest.

Furthermore, to assess potential correlations between the three dependent variables a Spearman correlation analysis was conducted. Spearman’s rank correlation coefficient ranges from -1 (perfect negative relationship) to 1 (perfect positive relationship), with 0 indicating no relationship. Significances were sought at the p < 0.05 level, and the p-values are represented as: *p < 0.05, **p < 0.01 and ***p < 0.001. Unless otherwise specified, data are presented as mean ± standard deviation (SD).

## RESULTS

### Spatial sensitivity of the palm surface

The sensitivity to mechanical stimuli was highly heterogeneous across the palm surface (Fig. 2 and Table S1). The distal phalanges (*i*.*e*., fingertips) of the 1^st^ and 2^nd^ fingers exhibited the lowest detection thresholds (0.03 ± 0.02 g), and in contrast, the highest pain thresholds (96.0 ± 95.8 g and 95.5 ± 95.7 g, respectively). Conversely, proximal areas of the hand such as the wrist displayed high tactile threshold and low pain thresholds (0.10 ± 0.06 g for detection and 34.7 ± 58.9 for pain). The intensity of evoked pain showed a similar pattern, with the 1^st^ and 2^nd^ fingers exhibiting lower scores (3.3 ± 2.1 g and 3.3 ± 2.3 g, respectively), while the wrist reported higher pain values (4.1 ± 2.4 au).

**Figure 2.**
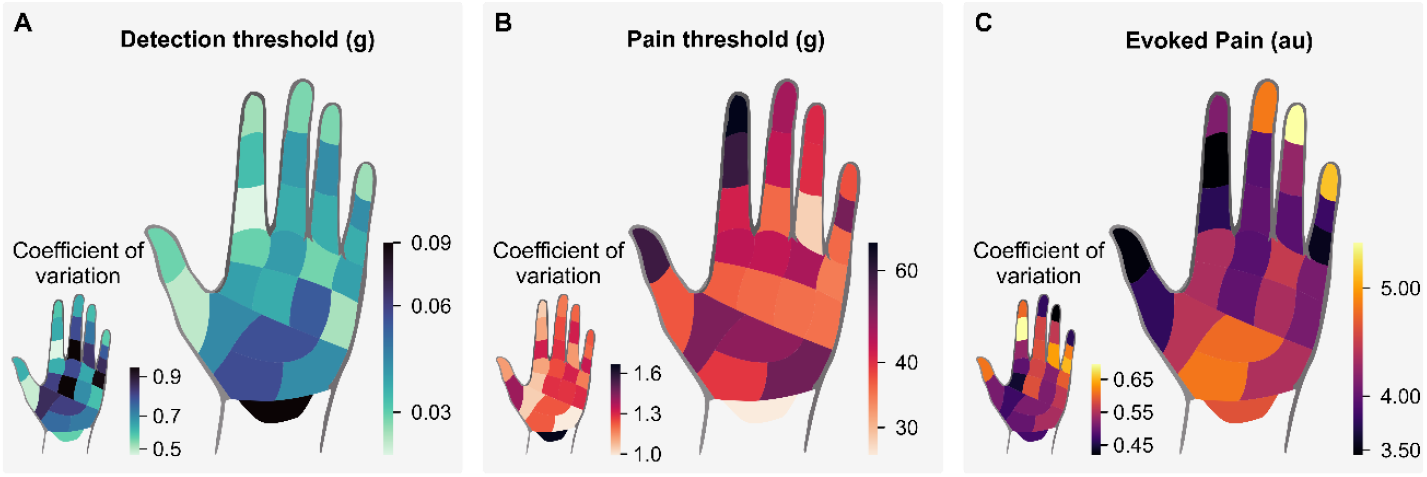
Spatial sensitivity mapping of the palm surface. Heatmaps illustrating mechanical sensitivity across 27 distinct areas on the palm. (A) Detection Threshold: quantified in grams (g) on a logarithmic scale. (B) Pain Threshold: quantified in grams (g) on a logarithmic scale. (C) Evoked Pain: assessed using a Numeric Rating Scale (NRS) from 0 to 10, reported in arbitrary units (au).

### Correlation across areas of detection and pain thresholds, and evoked pain

The inverse relationship between detection and pain sensitivities found at the fingertips motivated us to test for possible correlations between all 3 measures across the 27 areas of the hand. This analysis revealed an inverse relationship between tactile and pain thresholds (Fig. 3A), but not between tactile thresholds and evoked pain (Fig. 3B). Both pain measures were strongly, though not perfectly correlated (Fig. 3C).

**Figure 3.**
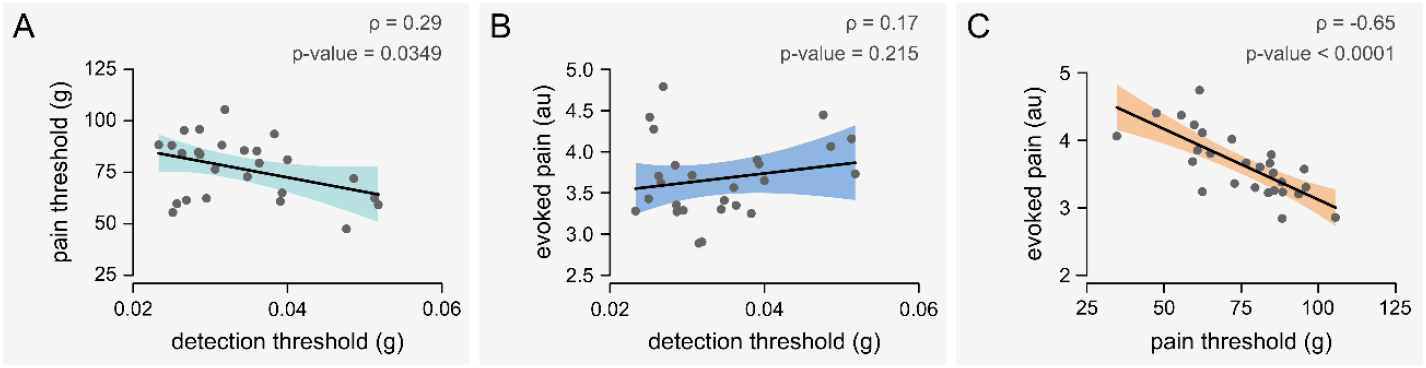
Relationship between sensitivities. (A–C) Each point represents the mean of an area (n = 27), averaging the measurements obtained in all the hands assessed (n = 66). Scatterplots show individual points along with the regression line ± SD. Relationships between variables were assessed using Spearman correlation, and each panel reports Spearman’s rank correlation coefficient (ρ) and p-value.

### Finger and phalanx sensitivity

Next, and due to their relevance in tactile exploration and object manipulation, we compared the sensitivities of the five fingers of the hand and their respective phalanges. The 1^st^ and 2^nd^ stood out with notably lower detection thresholds, higher pain thresholds, and lower evoked pain scores (Fig. 4A-C). I.e. they are very sensitive to touch, but relatively insensitive to pain. When each phalanx was analyzed separately, the middle phalanx displayed significantly higher tactile and pain thresholds, while the distal phalanx was the most sensitive (Fig. 4D-F).

**Figure 4.**
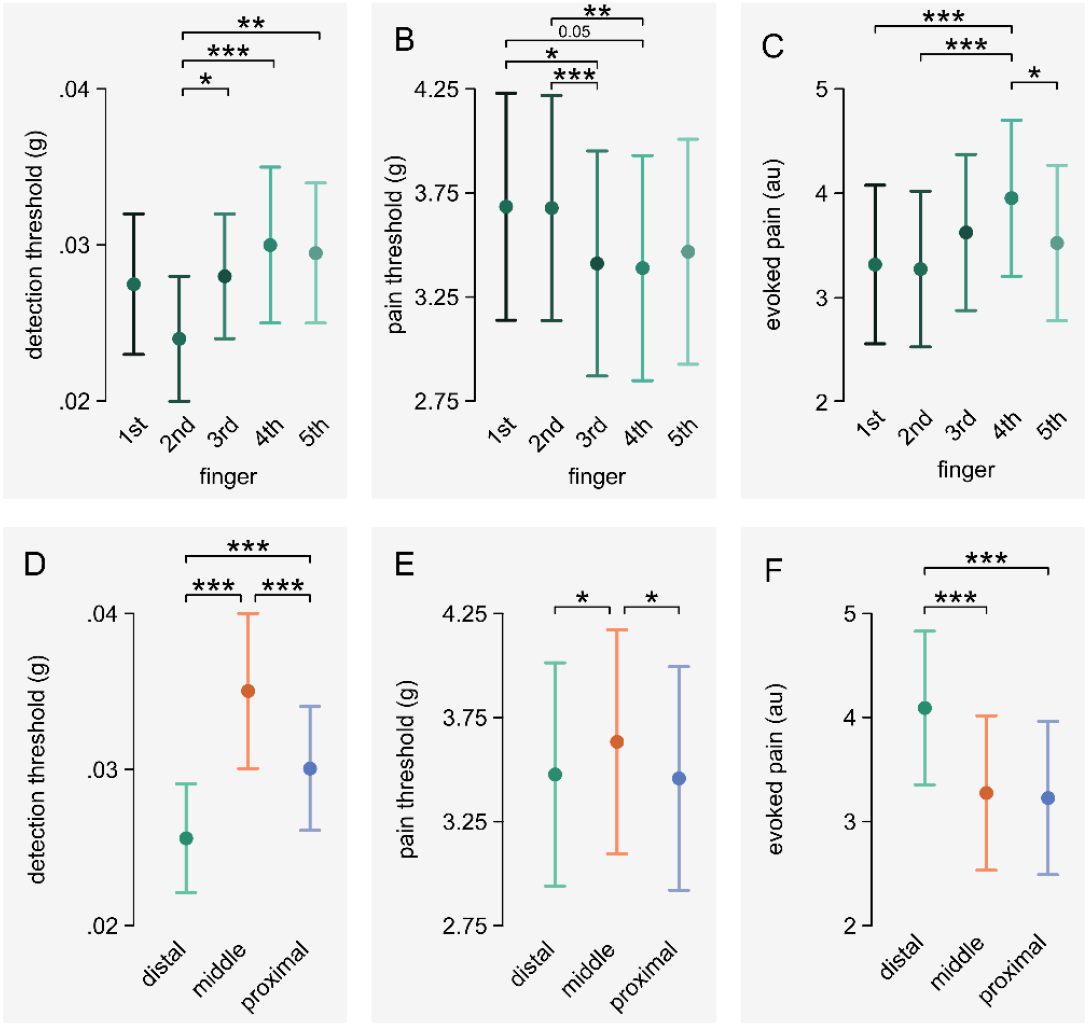
Finger and phalanx sensitivity. Mean values across individual fingers (A-C) and phalanges (D-F). Detection and pain thresholds are quantified in grams (g) and Evoked Pain in arbitrary units (au). Data are estimated marginal means (EMMs) from the models, shown as mean ± 95% CI, with significance levels denoted as follows: *p < 0.05, **p < 0.01, ***p < 0.001. The analysis for detection threshold utilized a Generalized Mixed Model, while Linear Mixed Models were applied for pain threshold and evoked pain. In all cases, a post-hoc test was conducted with Holm correction for multiple comparisons.

### Relations between mechanical sensitivity and hand dominance and sex

We also explored the effect of hand dominance in mechanical sensitivity. The dominant hand exhibited lower sensitivity compared to the non-dominant (i.e., higher tactile threshold and lower evoked pain) (Fig. 5A). To confirm that this was attributable to the dominance of the hand and not to laterality (right vs. left), we further compared the left hand against the right hand of each subject, segregating the analysis based on handedness. As anticipated, side differences were inverted between groups (i.e., less sensitivity in the left hand for left-handers and the right hand for right-handers), indicating a clear link to hand dominance.

**Figure 5.**
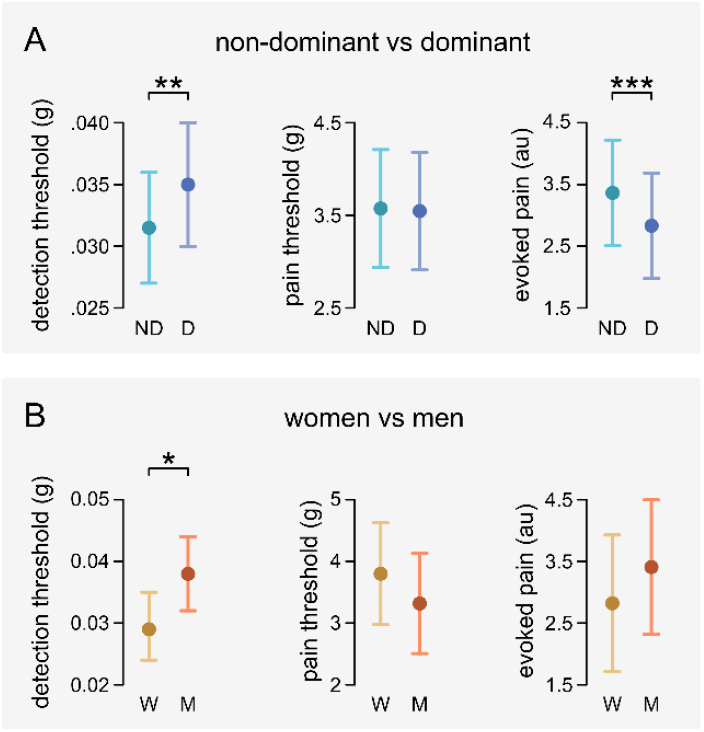
Sensitivity differences for hand dominance and sex. Comparison between non-dominant and dominant hands (A) and between males and females (B). Data are EMMs from the models, shown as mean ± 95% CI, with significance levels denoted as follows: *p < 0.05, **p < 0.01, ***p < 0.001. The analysis for detection threshold utilized a Generalized Mixed Model, while Linear Mixed Models were applied for pain threshold and evoked pain.

Finally, we investigated whether sex might also influence sensitivity (Venkatesan *et al*., 2015). Females exhibited lower detection thresholds compared to males, while no significant differences were observed in pain-related variables (Fig. 5B).

## DISCUSSION

Our results reveal a coarse-grained spatial distribution of mechanical sensitivity across 27 areas of the palm of the human hand. Tactile sensitivity was negatively correlated with pain sensitivity, and differences in mechanical sensitivity were found between dominant and non-dominant hands as well as between males and females.

### Spatial organization of tactile sensitivity

Regarding tactile sensitivity, the areas with lower detection thresholds were the fingertips, in particular of the 1^st^ and 2^nd^ digits, consistent with previous results (Craig & Lyle, 2001; Bowden & McNulty, 2013*b*), and supporting our departing hypothesis. But what determines the tactile sensitivity of each area? While different skin thickness might provide for an explanation, Lundström *et al*., 2018 found that skin thickness on the index finger does not notably affect perception of temperature or vibration. In contrast, Güçlü *et al*., 2008 did find higher Merkel-cell (SA1) densities in fingers 1^st^ and 2^nd^ than in the 4^th^ and another study showed that afferent receptors in the hand are densely packed in the distal ends of the fingertips and much less so in the palm (Corniani & Saal, 2020). Moreover, non-neural anatomical factors must be considered, such as variations in the arrangement of muscles, tendons, fascia, and blood vessels. In this line, Hudson *et al*., (2015) showed that increasing fingertip compliance via venous occlusion alters cutaneous afferent firing.

### Tactile and pain sensitivity are inversely correlated

To our surprise, mechanical detection thresholds were negatively correlated with pain thresholds along the whole palmar surface. As an example, the first two fingers required the smaller forces for stimulus detection, but the greatest forces to generate pain.

From a functional perspective, the first two fingers presenting increased tactile and reduced pain sensitivity may optimize object manipulation and exploration, as this increases the dynamic range of pressures these fingers can detect without triggering protective responses. This is particularly useful, as these fingers perform the pincer grasp (the precise grasping of an object between the thumb and forefinger), one of the most essential human sensorimotor skills (Napier, 1956).

Although purely speculative, the physiological substrate of our results might involve both peripheral and central mechanisms. Mechanical detection thresholds are mediated by low-threshold mechanoreceptors and pain thresholds by polymodal and mechano-nociceptors (Lumpkin & Caterina, 2007). It is intriguing therefore that activation of such independent receptors results in inversely correlated responses.

It is well-documented that increased usage of peripheral tissues stimulates the growth of epidermal cells, resulting in a thicker epidermis (Suominen *et al*., 1978; Biggs *et al*., 2020; Chien & Tsai, 2023). It could also be the case that a high density of low-threshold mechanoreceptors might be associated with a lower abundance of mechanosensitive nociceptors, and *vice versa*. However, characterization of the nociceptive innervation of the hand remains incomplete in comparison to tactile receptors (Johansson & Vallbo, 1979). In this line, Nolano *et al*., 2003 provided a histological characterization of the glabrous skin, revealing that the fingertips possess a complex and irregular nociceptive innervation that is relatively sparse. However, their analysis was restricted to the fingertips. Consequently, a map of the hand is required to test whether the opposing tactile and pain sensitivity patterns reported here are driven by peripheral afferents innervation density.

Previous works reported that frequent use and training can improve tactile acuity (Wong *et al*., 2013). This is proposed to be caused by central adaptation mechanisms such as dynamic intracortical inhibition, which enhances sensory contrast by suppressing widespread or redundant neural activity, thereby increasing the signal-to-noise ratio and allowing for more precise stimuli processing (Popescu *et al*., 2010, 2013; Venkatesan *et al*., 2014). Furthermore, it is well-known that sensory modalities sensed separately in the periphery converge in the central nervous system, leading to cross-talk and interaction (Melzack & Wall, 1965; Ma, 2010; Prescott *et al*., 2014; Velasco *et al*., 2024). These crosstalk mechanisms could explain the inverse relationship between detection and pain thresholds, as a mechanical stimulus delivered to the fingertips would evoke high levels of neural activity in the low-threshold mechanoreceptors pathway that may then inhibit nociceptive processing, hindering the perception of pain. Otherwise, when delivered to an area with fewer low-threshold mechanoreceptors (or in which those are activated less), nociceptive activation would be unrestricted to cause pain.

### Relation between sensitivity and hand dominance and sex

The dominant hand of the subjects was less sensitive to tactile stimuli than its non-dominant counterpart, irrespective of them being left- or right-handed. While pain thresholds were not different, the intensity of pain produced in response to a standardized painful stimulus was also reduced in the dominant hand.

Adaptations derived from usage may contribute to explaining these differences. A thicker skin in the dominant hand induced by mechanical stress may account for both decreased tactile sensitivity and diminished evoked pain (Vijusha & Ponmathi, 2018; Chien & Tsai, 2023). However, as previously noted, general skin thickness does not appear to dictate sensitivity (Lundström *et al*., 2018). Thus, this effect might be restricted to specific high-load sites rather than acting as a global dampening mechanism. Other potential contributors might be the perceptual habituation derived from frequent usage (Popescu *et al*., 2010; Venkatesan *et al*., 2014), as well as genetic factors, as sensitivity to touch and pain exhibits significant heritability (Fillingim *et al*., 2008; Frenzel *et al*., 2012). For instance, hand development is governed by genetic organizers and morphogens, such as SOXs and BMPs, which determine not only structural formation but also cutaneous receptor density (Jenkins & Lumpkin, 2017). These genetic factors, alongside the hypothesized X-linkage of handedness (Llaurens *et al*., 2009), could result in lateralized variations in sensitivity. Given that consistent hand preference typically emerges later in infancy (Hildreth, 1949), future longitudinal studies should investigate whether these sensory asymmetries are present from birth or are developed later in life.

Here we found that males showed reduced tactile sensitivity compared to females. This may be related to females presenting higher elasticity and reduced thickness in their skin, thus leading to greater indentations in response to application of the same force (Leveque *et al*., 1980; Cua *et al*., 1990; Sandby-Møller *et al*., 2003). Furthermore, female hands are overall smaller, and humans with smaller fingers have better passive tactile spatial perception, a phenomenon attributed to their higher innervation density (Peters *et al*., 2009; Peters & Goldreich, 2013). Contrasting to tactile thresholds, we found no significant differences in pain-related measures, in disagreement with several studies reporting interactions between sex and pain perception (Bartley & Fillingim, 2013), but agreeing with others reporting no differences between sexes (Racine *et al*., 2012).

### Study limitations

While the type of von Frey filaments used in this study are standard in clinical practice, it is worth noting several caveats when using them to measure mechanical thresholds. The non-linear increase in force from one Von Frey filament to the next can pose a challenge in accurately determining the threshold. In addition, thinnest von Frey filament produces a force of 0.02 g. But this filament was detected by most subjects, especially in the distal phalanx of the first and second fingers (in which 32 out of 33 subjects presented a threshold of 0.02 g). Therefore, the real detection limit in these areas remains undefined (though below 0.02 g), and this caused an artificial ‘floor effect’ in the distribution of the data in these areas, with most data points grouped in the minimal value. Furthermore, increasing the proportion of left-handed subjects would have been desirable, as the limited number of left-handed participants (7 out of 33) in our study hinders deeper analysis related to handedness. Additionally, our young sample (mean age 26) limits generalizability to older populations. In future studies, a comprehensive evaluation and combination of results from various stimulation modalities (mechanical, thermal and chemical), as well as comparisons between healthy and pathological subjects, would yield further useful knowledge to this surprisingly unexplored field.

## CONCLUSION

Tactile sensitivity is heterogeneously distributed across the palm of the human hand, which is partly explained by its innervation profile. This is also the case for pain sensitivity, though the underlying mechanisms are less characterized. Interestingly, tactile and pain sensitivity are inversely correlated. Areas with high tactile sensitivity present lower pain sensitivity, making them more suited for object exploration and manipulation, of which the most notable examples are the fingertips of the first two fingers. Conversely, areas with high pain sensitivity present less tactile acuity, displaying a more protective profile, as is the case of the wrist. Furthermore, the mechanical sensitivity relates to both the hand dominance and the sex: dominant hands are less sensitive to both pain and touch, and women have higher tactile sensitivity than men. These findings contribute to a better understanding of touch and pain and the physiology of the human hand.

## COMPETING INTERESTS

No competing interests to declare.

## FUNDING

No funding was received for this study.

## ACKNOWLEDGEMENTS

We acknowledge the Francisco Javier Ortega Physiotherapy Clinic for the facilities and material contribution (Elche, Alicante, Spain). We thank the study volunteers, who generously dedicated their time and effort to this study.

## AUTHOR’S CONTRIBUTION

Conceptualization and design of the study, V.M., F.A.C., M.D.M., E.V.; acquisition, analysis or interpretation of data for the work: V.M, M.J.G, M.M, E.V.; drafting the work or revising it critically for intellectual content: V.M, M.J.G, M.M., M.D.M., K.T. E.V; figures F.A.C and V.M. All authors have approved the final version of the manuscript submitted for publication and agree to be accountable for all aspects of this work in ensuring that questions related to the accuracy or integrity of any part of the work are appropriately investigated and resolved. All people designated as authors qualify for authorship and all those who qualify for authorship are listed.

---

**Table S1.** Spatial sensitivity mapping of the palm surface. Thresholds are expressed in grams (g) and evoked pain scores in arbitrary units from 0-10.

| Area | Detection threshold<br>(Mean ± SD) | Pain threshold<br>(Mean ± SD) | Evoked pain score<br>(Mean ± SD) |
| --- | --- | --- | --- |
| 1 | 0.06 ± 0.04 | 39.34 ± 48.15 | 4.82 ± 2.4 |
| 2 | 0.06 ± 0.05 | 48.49 ± 59.62 | 4.77 ± 2.45 |
| 3 | 0.04 ± 0.03 | 51.9 ± 51.37 | 4.37 ± 2.37 |
| 4 | 0.09 ± 0.05 | 26.52 ± 44.2 | 4.64 ± 2.21 |
| 11 | 0.03 ± 0.02 | 58.45 ± 65.82 | 3.49 ± 2.13 |
| 12 | 0.02 ± 0.01 | 36.95 ± 51.82 | 3.77 ± 1.92 |
| 13 | 0.04 ± 0.04 | 50.39 ± 52.42 | 4.41 ± 2.03 |
| 21 | 0.03 ± 0.02 | 67.98 ± 71.21 | 4.14 ± 2.49 |
| 22 | 0.03 ± 0.02 | 59.1 ± 65.61 | 3.45 ± 2.4 |
| 23 | 0.02 ± 0.01 | 41.75 ± 55.03 | 3.74 ± 1.98 |
| 24 | 0.03 ± 0.02 | 43.76 ± 48.66 | 4.36 ± 2.15 |
| 25 | 0.04 ± 0.03 | 36.42 ± 38.38 | 4.36 ± 1.92 |
| 31 | 0.03 ± 0.02 | 46.36 ± 54.8 | 4.84 ± 2.25 |
| 32 | 0.04 ± 0.03 | 43.42 ± 54.09 | 3.95 ± 2.26 |
| 33 | 0.04 ± 0.03 | 35.59 ± 45.76 | 3.99 ± 2.33 |
| 34 | 0.03 ± 0.03 | 38.28 ± 46.56 | 4.51 ± 2.62 |
| 35 | 0.05 ± 0.04 | 34.76 ± 50.66 | 5.23 ± 2.59 |
| 41 | 0.03 ± 0.02 | 40.64 ± 49.25 | 5.42 ± 2.27 |
| 42 | 0.04 ± 0.03 | 40.31 ± 53.16 | 4.24 ± 2.37 |
| 43 | 0.04 ± 0.03 | 28.34 ± 32.05 | 3.95 ± 2.52 |
| 44 | 0.03 ± 0.02 | 45.71 ± 59.95 | 4.48 ± 2.32 |
| 45 | 0.05 ± 0.04 | 35.25 ± 45.1 | 4.34 ± 2.23 |
| 51 | 0.03 ± 0.02 | 37.48 ± 46.03 | 5.14 ± 2.37 |
| 52 | 0.04 ± 0.03 | 51.15 ± 61.63 | 3.82 ± 2.35 |
| 53 | 0.04 ± 0.03 | 35.65 ± 47.51 | 3.55 ± 2.31 |
| 54 | 0.04 ± 0.04 | 34.35 ± 40.35 | 4.11 ± 2.44 |
| 55 | 0.03 ± 0.02 | 35.15 ± 45.53 | 4.1 ± 2.29 |

## Notes

### Competing Interest Statement

The authors have declared no competing interest.

## REFERENCES

Abraira VE & Ginty DD (2013). The sensory neurons of touch. Neuron 79, 618–639.

Ackerley R, Saar K, McGlone F & Wasling HB (2014). Quantifying the sensory and emotional perception of touch: differences between glabrous and hairy skin. Frontiers in behavioral neuroscience; DOI: 10.3389/FNBEH.2014.00034.

Bartley EJ & Fillingim RB (2013). Sex differences in pain: a brief review of clinical and experimental findings. BJA: British Journal of Anaesthesia 111, 52.

Beltrá P, Ruiz-Del-Portal I, Ortega FJ, Valdesuso R, Delicado-Miralles M & Velasco E (2022). Sensorimotor effects of plasticity-inducing percutaneous peripheral nerve stimulation protocols: a blinded, randomized clinical trial. Eur J Pain 26, 1039–1055.

Biggs LC, Kim CS, Miroshnikova YA & Wickström SA (2020). Mechanical Forces in the Skin: Roles in Tissue Architecture, Stability, and Function. Journal of Investigative Dermatology 140, 284–290.

Bowden JL & McNulty PA (2013a). The magnitude and rate of reduction in strength, dexterity and sensation in the human hand vary with ageing. Experimental Gerontology 48, 756–765.

Bowden JL & McNulty PA (2013b). Age-related changes in cutaneous sensation in the healthy human hand. Age (Dordr) 35, 1077–1089.

Catley MJ, O’Connell NE, Berryman C, Ayhan FF & Moseley GL (2014). Is tactile acuity altered in people with chronic pain? a systematic review and meta-analysis. The journal of pain 15, 985–1000.

Chien C-W, Bagraith KS, Khan A, Deen M & Strong J (2013). Comparative Responsiveness of Verbal and Numerical Rating Scales to Measure Pain Intensity in Patients With Chronic Pain. The Journal of Pain 14, 1653–1662.

Chien WC & Tsai TF (2023). The Pressurized Skin: A Review on the Pathological Effect of Mechanical Pressure on the Skin from the Cellular Perspective. International Journal of Molecular Sciences 24, 15207.

Cobo R, García-Piqueras J, Cobo J & Vega JA (2021). The Human Cutaneous Sensory Corpuscles: An Update. Journal of Clinical Medicine 10, 1–12.

Cole K, Rotella D & Harper J (1998). Tactile impairments cannot explain the effect of age on a grasp and lift task. Experimental brain research Experimentelle Hirnforschung Expérimentation cérébrale 121, 263–269.

Corniani G & Saal HP (2020). Tactile innervation densities across the whole body. Journal of neurophysiology 124, 1229–1240.

Craig JC & Lyle KB (2001). A comparison of tactile spatial sensitivity on the palm and fingerpad. Perception & psychophysics 63, 337–347.

Cua AB, Wilhelm KP & Maibach HI (1990). Frictional properties of human skin: relation to age, sex and anatomical region, stratum corneum hydration and transepidermal water loss. Br J Dermatol 123, 473–479.

Farrell MJ, Gibson SJ, McMeeken JM & Helme RD (2000). Increased movement pain in osteoarthritis of the hands is associated with Aβ-mediated cutaneous mechanical sensitivity. The Journal of Pain 1, 229–242.

Fillingim RB, Wallace MR, Herbstman DM, Ribeiro-Dasilva M & Staud R (2008). Genetic contributions to pain: a review of findings in humans. Oral diseases 14, 673.

Frenzel H, Bohlender J, Pinsker K, Wohlleben B, Tank J, Lechner SG, Schiska D, Jaijo T, Rüschendorf F, Saar K, Jordan J, Millán JM, Gross M & Lewin GR (2012). A Genetic Basis for Mechanosensory Traits in Humans. PLoS Biology 10, 1001318.

Gellis M & Pool R (1977). Two-point discrimination distances in the normal hand and forearm: application to various methods of fingertip reconstruction. Plast Reconstr Surg 59, 57–63.

Güçlü B, Mahoney GK, Pawson LJ, Pack AK, Smith RL & Bolanowski SJ (2008). Localization of Merkel cells in the monkey skin: an anatomical model. Somatosensory & motor research 25, 123–138.

Hildreth G (1949). The development and training of hand dominance; characteristics of handedness. J Genet Psychol 75, 197–220.

Hudson KM, Condon M, Ackerley R, McGlone F, Olausson H, Macefield VG & Birznieks I (2015). Effects of changing skin mechanics on the differential sensitivity to surface compliance by tactile afferents in the human finger pad. Journal of Neurophysiology 114, 2249–2257.

Jarocka E, Pruszynski JA & Johansson RS (2021). Human touch receptors are sensitive to spatial details on the scale of single fingerprint ridges. Journal of Neuroscience 41, 3622–3634.

Jenkins BA & Lumpkin EA (2017). Developing a sense of touch. Development 144, 4078–4090.

Johansson RS & Vallbo AB (1979). Tactile sensibility in the human hand: relative and absolute densities of four types of mechanoreceptive units in glabrous skin. J Physiol 286, 283–300.

Kalisch T, Tegenthoff M & Dinse HR (2008). Improvement of sensorimotor functions in old age by passive sensory stimulation. Clin Interv Aging 3, 673–690.

Leveque JL, de Rigal J, Agache PG & Monneur C (1980). Influence of ageing on the in vivo extensibility of human skin at a low stress. Arch Dermatol Res 269, 127–135.

Llaurens V, Raymond M & Faurie C (2009). Why are some people left-handed? An evolutionary perspective. Philosophical transactions of the Royal Society of London Series B, Biological sciences 364, 881–894.

Lumpkin EA & Caterina MJ (2007). Mechanisms of sensory transduction in the skin. Nature 445, 858–865.

Lundström R, Dahlqvist H, Hagberg M & Nilsson T (2018). Vibrotactile and thermal perception and its relation to finger skin thickness. Clinical neurophysiology practice 3, 33–39.

Ma Q (2010). Labeled lines meet and talk: population coding of somatic sensations. J Clin Invest 120, 3773–3778.

McIntyre S, Nagi SS, McGlone F & Olausson H (2021). The Effects of Ageing on Tactile Function in Humans. Neuroscience 464, 53–58.

Melzack R & Wall PD (1965). Pain mechanisms: a new theory. Science 150, 971–979.

Napier J (1962). The evolution of the hand. Sci Am 207, 56–62.

Napier JR (1956). The prehensile movements of the human hand. J Bone Joint Surg Br 38-B, 902–913.

Nolano M, Provitera V, Crisci C, Stancanelli A, Wendelschafer-Crabb G, Kennedy WR & Santoro L (2003). Quantification of myelinated endings and mechanoreceptors in human digital skin. Ann Neurol 54, 197–205.

Novak CB, Kelly L & Mackinnon SE (1992). Sensory recovery after median nerve grafting. The Journal of hand surgery 17, 59–68.

Peters RM & Goldreich D (2013). Tactile Spatial Acuity in Childhood: Effects of Age and Fingertip Size. PLoS ONE 8, 84650.

Peters RM, Hackeman E & Goldreich D (2009). Diminutive Digits Discern Delicate Details: Fingertip Size and the Sex Difference in Tactile Spatial Acuity. J Neurosci 29, 15756–15761.

Popescu EA, Barlow SM, Venkatesan L, Wang J & Popescu M (2013). Adaptive changes in the neuromagnetic response of the primary and association somatosensory areas following repetitive tactile hand stimulation in humans. Human Brain Mapping 34, 1415–1426.

Popescu M, Barlow S, Popescu EA, Estep ME, Venkatesan L, Auer ET & Brooks WM (2010). Cutaneous stimulation of the digits and lips evokes responses with different adaptation patterns in primary somatosensory cortex. NeuroImage 52, 1477–1486.

Prescott SA, M. Q & De Koninck Y (2014). Normal and abnormal coding of somatosensory stimuli causing pain. Nat Neurosci 17, 183–191.

Racine M, Tousignant-Laflamme Y, Kloda LA, Dion D, Dupuis G & Choinière M (2012). A systematic literature review of 10 years of research on sex/gender and experimental pain perception - part 1: are there really differences between women and men? Pain 153, 602–618.

Sandby-Møller J, Poulsen T & Wulf HC (2003). Epidermal thickness at different body sites: relationship to age, gender, pigmentation, blood content, skin type and smoking habits. Acta dermato-venereologica 83, 410–413.

Suominen H, Heikkinen E, Moisio H & Viljamaa K (1978). Physical and chemical properties of skin in habitually trained and sedentary men. Br J Dermatol 99, 147–154.

Tremblay F, Mireault A-C, Dessureault L, Manning H & Sveistrup H (2005). Postural stabilization from fingertip contact. Exp Brain Res 164, 155–164.

Velasco E, Zaforas M, Acosta MC, Gallar J & Aguilar J (2024). Ocular surface information seen from the somatosensory thalamus and cortex. The Journal of PhysiologyJP285008.

Venkatesan L, Barlow SM & Kieweg D (2015). Age- and sex-related changes in vibrotactile sensitivity of hand and face in neurotypical adults. Somatosensory & motor research 32, 44–50.

Venkatesan L, Barlow SM, Popescu M & Popescu A (2014). Integrated approach for studying adaptation mechanisms in the human somatosensory cortical network. Experimental brain research 232, 3545–3554.

Vijusha T & Ponmathi P (2018). Comparison of Two Point Discrimination of Palmar Surface of Distal Phalanx of Thumb between Dominant and Non Dominant in Labourers. International Journal of Medical Science and Dental Research 01, 01–06.

Wong M, Peters RM & Goldreich D (2013). A physical constraint on perceptual learning: tactile spatial acuity improves with training to a limit set by finger size. The Journal of neuroscience : the official journal of the Society for Neuroscience 33, 9345–9352.

